# Metabolic homeostasis favors tolerance to persistent viral infection in *Aedes aegypti* mosquitoes

**DOI:** 10.1101/2025.07.23.666353

**Authors:** Hugo D. Perdomo, Ayda Khorramnejad, Francesco Chen, Alejandro N. Lozada-Chávez, Mariangela Bonizzoni

## Abstract

The mosquito *Aedes aegypti* harbors a wide range of persistent viral infections, including both medically important arboviruses and insect-specific viruses (ISVs). However, the mechanisms underpinning establishment and maintenance of persistent viral infections remain poorly understood despite them influencing viral dynamics and transmission. Differently than arboviruses, which circulate at lower frequencies, ISVs are highly prevalent in field mosquitoes, thus representing ideal models to study the mechanisms of persistence. On this basis, we followed infection of the ISV cell-fusing agent virus (CFAV) in two *Ae. aegypti* laboratory populations, Liverpool (LVP) and ZacPanda (ZacP), and studied their immune responses, including resistance and tolerance, along with physiological and fitness impacts of CFAV infection. Our results demonstrate that CFAV establishes life-long infections with a comparable activation of resistance mechanisms, in both LVP and ZacP mosquitoes. In contrast, ZacP mosquitoes exhibited greater tolerance to CFAV than LVP accompanied by the ability to maintain metabolic homeostasis during infection. In neither ZacP nor LVP, CFAV resulted in reproductive impairment. These findings highlight a key role for tolerance mechanisms, which through regulation of energetic resources, contribute to sustain persistent viral infections in mosquitoes. Our results have implications for viral transmission dynamics and vector control strategies.

## Introduction

When an acute viral infection is not cleared, a persistent infection develops with viral particles being produced for long periods of time [1]. Studies on persistent viral infections in mammals have identified three key elements of persistent infections [1]. First, the virus must evade or modulate the host immune response to establish a persistent infection. Second, the virus may employ unique replication strategies and regulate either cellular and/or viral expression to maintain a non-lytic infection. Third, continuous viral replication impairs cellular homeostasis. These processes may result in disease symptoms on the host, such as the case of the human immunodeficiency virus (HIV) or manifest as asymptomatic infections, as those caused by the lymphocytic choriomeningitis virus (LCMV) in mice [2]. Beyond mammals, the ability of some arthropod species to sustain viral replication for long periods of time enables them to efficiently transmit epidemiologically-relevant viruses as Dengue, Zika, Yellow Fever, West Nile and Chikungunya viruses, collectively called arboviruses [3, 4]. The mosquito *Aedes aegypti* and *Aedes albopictus* are the most important arboviral vectors around the world [5]. However, how persistent viral infections are reached and maintained in *Aedes* spp. mosquitoes is poorly understood.

Being invertebrates, *Aedes* spp. mosquitoes lack adaptive immunity and mostly rely on RNA interference (RNAi) mechanisms to fight viral infections [6]. RNAi comprises three independent, but interlinked, pathways named for their effector RNA molecules as small interfering (si)RNA, micro (mi)RNA and PIWI-interacting (pi)RNA pathways [7]. The siRNA pathway is activated when double-stranded RNA (dsRNA) molecules are detected and cleaved into 21 nucleotides (nt) long fragments by Dicer-2 (Dcr2) [8]. These fragments are then loaded into Argonaute-2 (Ago2) that degrades target single-stranded RNA molecules based on sequence complementarity [9]. siRNAs are found in mosquitoes as early as 2 days post infection (dpi) and up to 14 dpi [10, 11], indicating mosquitoes actively control viral replication during persistent infection. Mosquitoes were also shown to control viral replication in ovaries through piRNAs derived from endogenous viral elements (EVEs) [12] and through Dcr2-dependent generation of viral DNA fragments from infecting RNA viruses, which further feed the siRNA pathway [13, 14]. Importantly, knock-out of both *dcr2* and *ago2* not only leads to hyper-viral infection, showing their essential role in controlling viral replication, but also causes reduced mosquito longevity [15, 16]. Despite all mosquitoes being able to mount an active RNAi response, their transcriptional response to a viral infection and the resulting fitness costs may be different [17, 18]. The variability across populations in the overall responses to a viral challenge supports the hypothesis that persistent viral infection entails not only resistance, but also tolerance mechanisms whereby mosquitoes control the cost of infection without impeding viral replication [19]. The mechanisms and impact of tolerance during persistent viral infections in mosquitoes remain largely unexplored.

Separating resistance versus tolerance responses during an active infection is not trivial, because underlying mechanisms and/or molecular effectors can be linked. For instance, a genome wide association study in *Drosophila melanogaster* found that tolerance and resistance are not completely independent traits and identified some single nucleotide polymorphisms (SNPs) that linked both mechanisms [20]. The intricacy between tolerance and resistance can be seen also at a single gene level. For instance, in *Dr. melanogaster* different SNPs in the protease CG3066, which is involved in melanization, were shown to affect differently resistance or tolerance to diverse bacterial infections [21]. Plant pathologists developed in the ‘90 a framework to disentangle the contribution of resistance and tolerance responses during an active infection [22, 23]. This framework requires the generation of dose-response curves that allow to highlight changes in host health dependent on increasing viral infection doses [24–26]; The comparison of the dose-response curves of at least two groups of samples can show which one suffered less, or tolerated better, viral infection [19]. A dose-response curve analyses can be easily adapted to animals.

Extensive entomological surveys report that in areas with intense arboviral transmission such as Yogyakarta City (Indonesia), Puerto Rico, Merida (Mexico) and Singapore, arboviral prevalence maintains below 10% (0.12%, 0.8%, 7.7% and 6.9%, respectively in the sites cited above) [27–30]. In contrast, mosquitoes are found most-frequently infected with insect-specific viruses (ISVs) [31–33], which can establish persistent infections comparable in duration to those of arboviruses. For instance, cell-fusing agent virus (CFAV), the first and, so far, best-characterized ISV [34–39], was shown to persist in mosquitoes for up to 35 days, often at higher viral loads than those observed with arboviruses [24, 40]. These findings indicate that ISVs play a more significant ecological role in mosquito biology, resulting in a higher evolutionary impact, than arboviruses.

On this basis, we used CFAV to test the relative contribution of resistance and tolerance mechanisms in the establishment of persistent infection in *Ae. aegypti* mosquitoes. We compared CFAV infections in two laboratory populations, ZacPanda (ZacP) and Liverpool (LVP), which differ in their origin and pattern of viral integrations with only ZacP mosquitoes harboring in their genomes EVEs with sequence similarity to CFAV [41] to further highlight the relative contribution of the siRNA and piRNA pathways in mosquito response to CFAV infection.

## Results

### CFAV induces persistent lifelong infection in *Ae. aegypti*

We injected three to eight days old *Ae. aegypti* females of LVP and ZacP laboratory populations with 10^3.32^ focus forming units (FFU) of CFAV and we quantified viral load one, three, seven and 14 dpi in both carcasses and their corresponding ovaries (Fig 1 A, B). We detected CFAV in carcasses of both populations starting 1 dpi, with viral load reaching a plateau by 3 dpi in LVP and by 7 dpi in ZacP (Fig 1A, S1 Table). At plateau, viral load was not significantly different between populations (S1 Table), which is consistent with the detection of a similar number of virions in both LVP and ZacP mosquitoes 14 dpi (Fig 1C). The viral titter at the plateau is of the same order of magnitude as the number of genomes that were detected in wild-caught mosquitoes [38], suggesting that our infections closely resemble natural infections.

**Figure 1:**
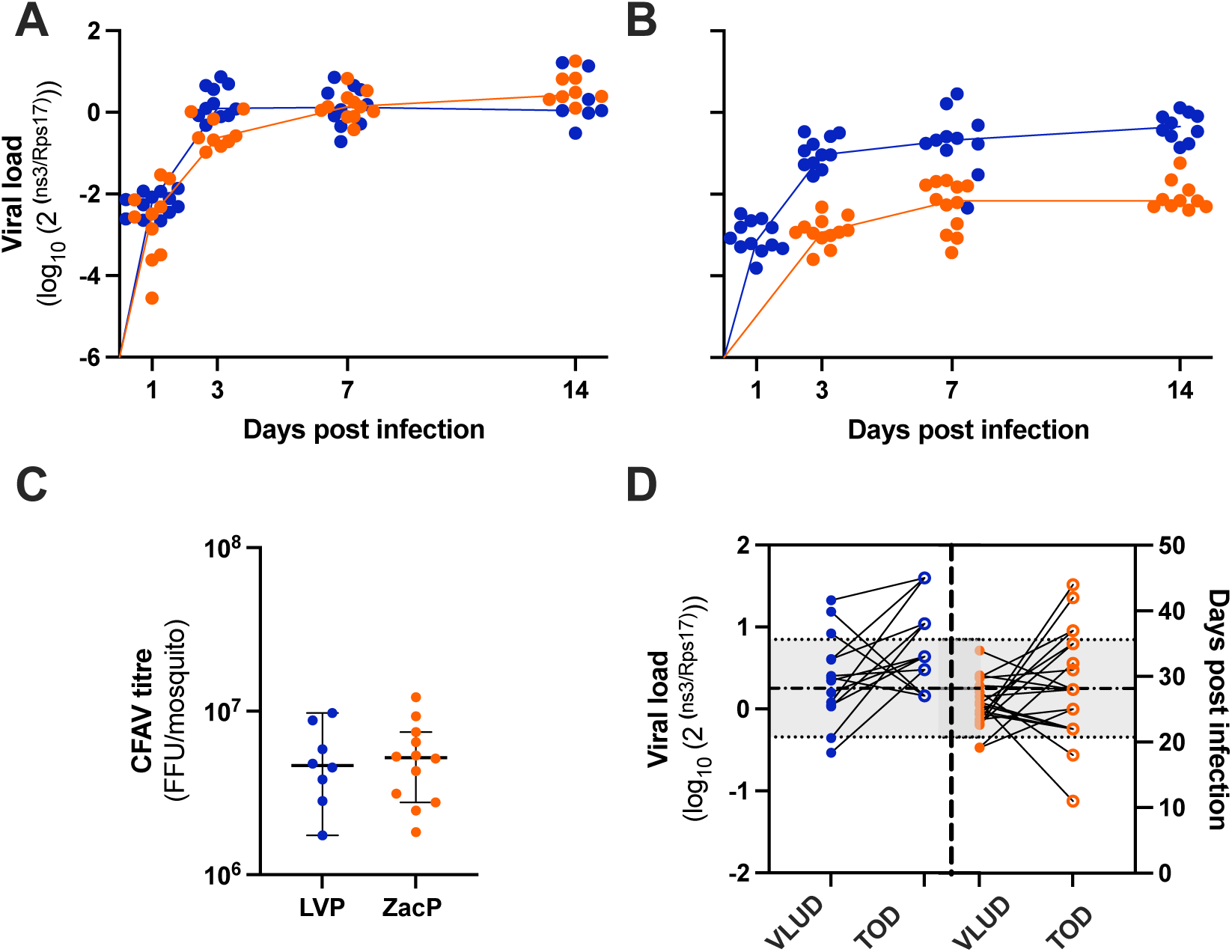
CFAV establishes lifelong persistent infection in *Ae. aegypti*. (**A**) quantification of CFAV viral genome load one, three, seven and 14 dpi in carcasses of LVP and ZacP *Ae. aegypti* mosquitoes. **(B)** quantification of CFAV viral genome load one, three, seven and 14 dpi in ovaries of LVP and ZacP *Ae. aegypti* mosquitoes. In both panels, the lines join the median viral load detected each day. **(C)** Quantification of viral particles by FFA 14 dpi. Each dot represents data from one sample, with indications of the median and 95% confidence intervals. **(D)** Quantification of viral load upon death (VLUD) as well as days elapsed after infection or time of death (TOD). A line was drawn to connect the values of VLUD and TOD of each single mosquito. Dash and dotted line represents the plateau in panel **A** and dotted lines represent the 95% confidence interval, respectively. Each dot represents data from an individual mosquito. In all panels data from LVP are in blue, those from ZacP in orange.

In contrast to carcasses, ovaries showed differences in CFAV infection between populations (Fig 1B, S1 Table). We detected CFAV in ovaries of LVP mosquitoes 1 dpi, with viral load reaching a plateau by 3dpi. We first observed CFAV in ovaries of ZacP mosquitoes 3 dpi and by 5 dpi viral load reached a plateau (time p < 0.0001), which was significantly lower than that of ovaries of LVP mosquitoes (Fig 1B, population p<0.0001). Detection of CFAV in ovaries prompted us to test for vertical transmission, which has been shown at variable frequency depending on mosquito populations [37, 39]. We observed vertical transmission only in LVP mosquitoes at a 1.59% frequency (S1 Fig.).

To verify if CFAV infection is maintained beyond 14 dpi, we quantified CFAV at the time of mosquito death (Fig 1D). We detected CFAV in every dead mosquito that had been injected with 10^3.32^ FFU of CFAV, proving injection always results in infection, which persists throughout the entire mosquito lifespan. We also observed that the viral load upon death (VLUD) is not significantly different than that at plateau [42] and does not correlate with the time of mosquito death (Fig 1D).

### Dose response curve analysis reveals lower tolerance in LVP than ZacP mosquitoes following CFAV infection

The observation that by 3-7 dpi CFAV load reaches a plateau, which is maintained until mosquito death (Fig 1A, D), support the conclusions that CFAV has limited virulence on *Ae. aegypti* mosquitoes and that mosquitoes manage the infection. To discriminate between resistance and tolerance, we applied a dose-response curve analysis followed by viral quantification at the time of mosquito death [22]. We infected mosquitoes with media and five increasing CFAV concentrations (10^1.32^ FFU, 10^2.32^ FFU, 10^3.32^ FFU, 10^4.32^ FFU, 10^5.32^ FFU) and monitored their survival as a proxy for health. We saw that CFAV reduces mosquito longevity in comparison to media-injected mosquitoes, starting at 10^3.32^ FFU (Fig 2A, B). Next, we plotted the survival time of each mosquito in relation to the infecting dose and fitted a four-parameter logistic model with a least squares regression to determine the survival time of media-injected mosquitoes, called vigor [26]; the longevity of mosquitoes when infected with the highest CFAV load, called severity [26], and the effective concentration 50 (EC_50_), which describes the viral load reducing host vigor by half, called sensitivity [26]. To compare the curves of LVP and ZacP mosquitoes, we constrained their slope, which denotes the rate of loss of mosquito health, to −1, as previously done [25, 26]. We observed that neither vigor nor sensitivity were significantly different between populations (S1 Table). In contrast, LVP mosquitoes showed higher severity than ZacP ones, with a 34.2% reduction of their lifespan in comparison to a 18.3% reduction of ZacP mosquitoes, with respect to their vigor (Fig 2C). These results show that ZacP mosquitoes tolerate CFAV infection better than LVP ones.

**Figure 2:**
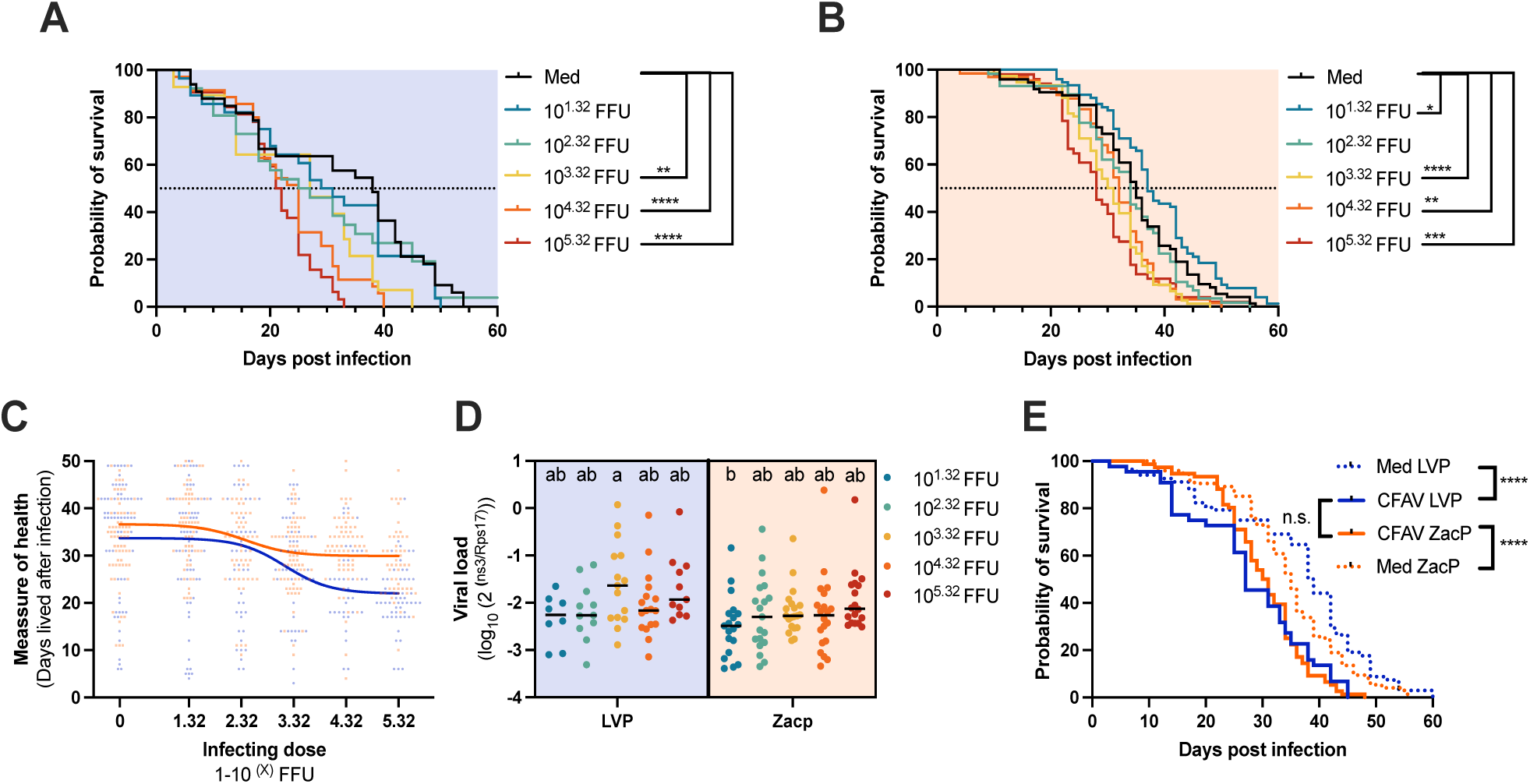
LVP mosquitoes have a lower tolerance to CFAV infection than ZacP. **(A)** probability of survival of LVP mosquitoes after infection with the five increasing CFAV concentrations and media. **(B)** Probability of survival of ZacP mosquitoes after infection with the five increasing CFAV concentrations and media. In both graphs, we used the Kaplan–Meier survival analysis with a cox proportional hazard test to determine the effect of CFAV infection on mosquito survival. **(C)** Tolerance curves were built by plotting infecting dose vs days lived after infection and a four-parameter logistic curve was fitted to the data. **(D)** Quantification of viral load of mosquitoes after their dead. Each dot represents a single mosquito; letters represent compact letter display of tukey’s multiple comparison test. **(E)** The probability of survival between not-infected mosquitoes and mosquitoes infected with 10^3.32^ viral particles, for both LVP and ZacP; significance was tested by the Kaplan–Meier survival analysis with a cox proportional hazard test. ns represents not significant, * P-value<0.05, ** P-value<0.01, *** P-value<0.001 and **** P-value<0.0001

Quantification of viral load at time of death showed that viral loads were similar regardless of the infecting doses or the populations. This result demonstrates that the observed difference in severity between ZacP and LVP mosquitoes is due to tolerance rather than resistance mechanisms. To avoid biases due to CFAV-dependent accelerated mortality of LVP mosquitoes, in all our following experiments, we injected mosquitoes with 10^3.32^ FFU, a dose that reduces mosquito longevity with respect to media infected mosquitoes, but results in no population differences in survival rates (Fig 2E).

### Presence of nrEVE-derived piRNAs does not influence CFAV load nor mosquito survival after infection

We compared resistance between ZacP and LVP mosquitoes by measuring gene expression of *ago2* and *dcr2* and performing small RNA sequencing of CFAV-infected mosquitoes 14 dpi, both in carcasses and ovaries.

We saw no difference in basal expression of *ago2* and *dcr2* between LVP and ZacP mosquitoes. After CAFV infection, we saw a reduced expression of *ago2* in both populations (Fig 3A), whereas expression of *dcr2* was reduced only in ZacP mosquitoes (Fig 3B). Despite the downregulation of *dcr2* and *ago2*, we saw an active siRNA response denoted by the presence of a sharp 21nt peak in both populations (Fig 3C). However, the size distribution of small RNAs (sRNAs) mapping to the CFAV genome was different between LVP and ZacP mosquitoes (Fig 3C). In LVP, we saw a peak of 21 nt sRNAs mapping to both the positive and negative strands and a smaller peak of 26-30 nt sRNAs mapping only to the positive strand (Fig 3C blue). We also observed uniform coverage of 21 nt sRNAs across the CFAV genome (Fig 3D blue) and mapping of 26-30 nt sRNAs to few spots (Fig 3E blue). In ZacP mosquitoes, we saw two peaks of 21nt and 26-30 nt sRNAs, which mapped to both strands (Fig 3C); 21 nt sRNAs showed a uniform coverage across the CFAV genome (Fig 3D orange) while 26-30 nt sRNAs mapped primarily to regions of the CFAV genome that correspond to EVE3 [41] and EVE2 [12], two CFAV-like integrations previously identified in the genome of ZacP mosquitoes and also mosquitoes from Thailand (Fig3E orange) [12, 41]. EVE3 and EVE2 are absent from the genome of LVP mosquitoes (Fig 3F). To test if the presence of these two CFAV-like nrEVEs influence the survival of ZacP mosquitoes following CFAV infection, we tested their prevalence in non-injected (NI), media injected and CFAV infected mosquitoes at 28 dpi. We did not observe differences in the prevalence of these nrEVEs across groups, suggesting that their presence is not crucial for survival (Fig 3G).

**Figure 3:**
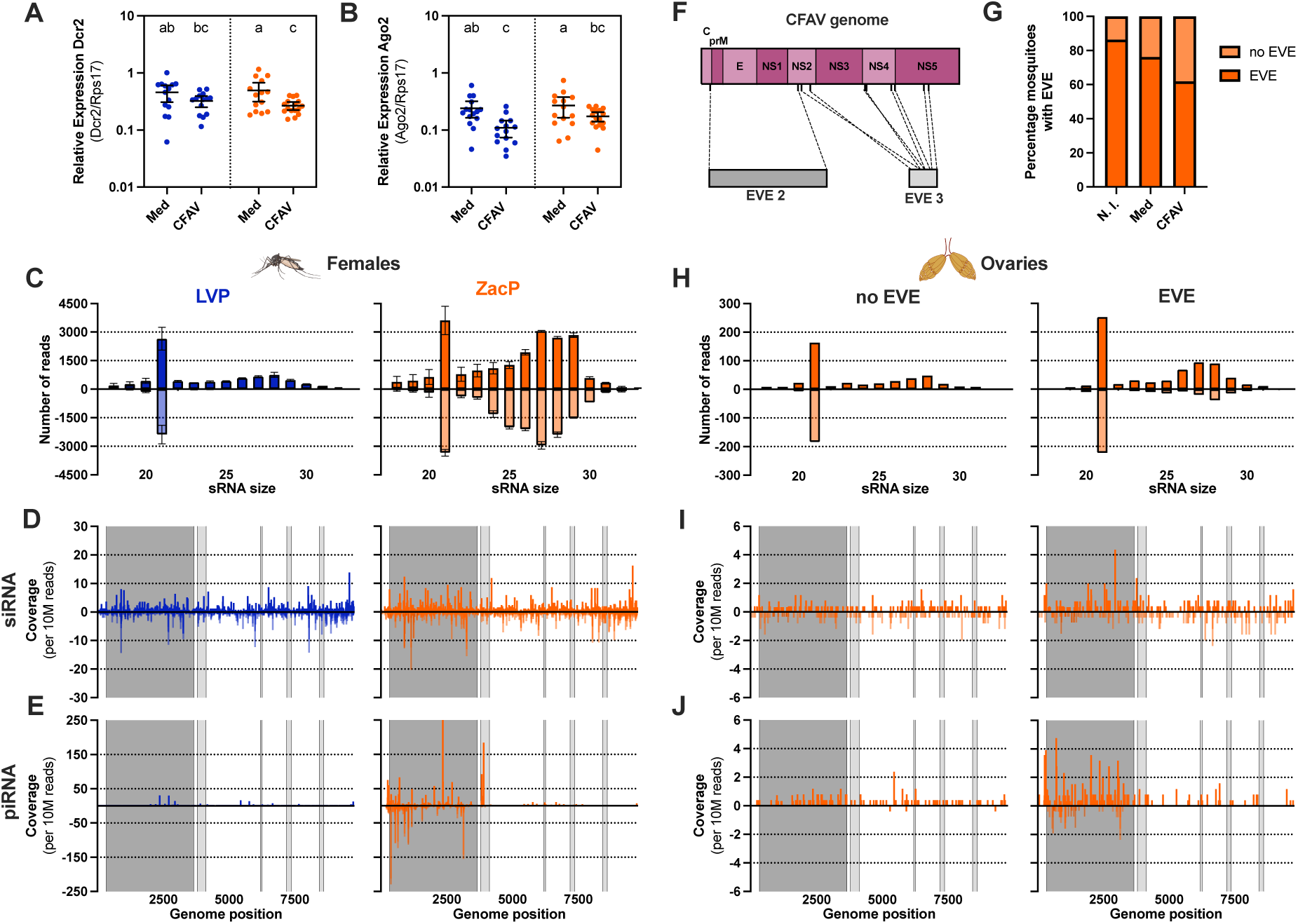
Comparable profile of siRNAs in ZacP and LVP, but nrEVEs-derived piRNAs only in ZacP mosquitoes. In all panels, data from LVP are in blue and from ZacP in orange. Expression levels of **(A)** *dcr2* and **(B)** *ago2* in LVP and ZacP mosquitoes 14 dpi with 10^3.32^ FFU of CFAV. Each points represents data from a single media injected (Med) or infected (CFAV) mosquitoes. Letters display results of the Tukey’s multiple comparison test. (**C**) Size distribution of sRNAs from LVP and ZacP whole mosquitoes mapping to the positive or negative strand of the CFAV genome. **(D)** Coverage of 21 nt reads across the CFAV genome. **(E)** Coverage of 26-30 nt reads across the CFAV genome. **(F)** Sketch of the CFAV genome showing its genes and the regions of similarity to EVE2 and EVE3. **(G)** Percentage of mosquitoes with EVE2 and EVE3 among not injected (NI), media injected (Media) and CFAV infected (CFAV) mosquitoes 28 dpi. (**H**) Size distribution of sRNAs from the ovaries of ZacP mosquito with or without an EVE mapping to the positive or negative strand of the CFAV genome. **(I)** Coverage of 21 nt reads across the CFAV genome. **(J)** Coverage of 26-30 nt reads across the CFAV genome. Dark grey and light grey shading represent areas of the CFAV genome with similarity to EVE2 and EVE3, respectively.

We further profiled small RNAs in ovaries because piRNAs derived from EVE2 were previously shown to contribute to control CFAV infection (Fig 3H-I) [12]. The size distribution of sRNAs showed that only ZacP mosquitoes with EVE2/3 had piRNAs with a negative strand polarity with respect to the CFAV genome (Fig 3H, S2 Fig). Presence of either EVE2 and/or EVE3 correlated with overall higher piRNAs abundance, with also presence of piRNAs mapping to the negative strand of the CFAV genome (Figure 3 J). Differently than piRNAs, siRNAs showed uniform coverage across the CFAV as observed in whole body and their load was not affected by the presence of EVEs (Fig 3I). Altogether, these results show that both LVP and ZacP mosquitoes mount an active RNAi response following CFAV infection and confirm production of nrEVE-derived piRNA-size sRNAs, in whole body and ovaries. While the presence of nrEVEs-derived piRNAs does not have a major effect on CFAV load in carcasses nor increase survival probability of infected mosquitoes, nrEVEs-derived piRNAs in ovaries of ZacP mosquitoes correlate with lower viral load with respect to ovaries of LVP mosquitoes.

### LVP mosquitoes have their metabolic homeostasis disrupted after CFAV infection

Having established that CFAV persistent infection requires both resistance and tolerance mechanisms and that ZacP mosquitoes are able to better sustain infection than LVP ones, independently from their ability to actively control CFAV replication, we investigated mechanisms of tolerance by comparing cellular homeostasis of the two laboratory populations. We measured carbohydrates, glycogen, proteins and lipids 14 days post CFAV infection in females of the two laboratory populations. (Fig 4A-D).

**Figure 4:**
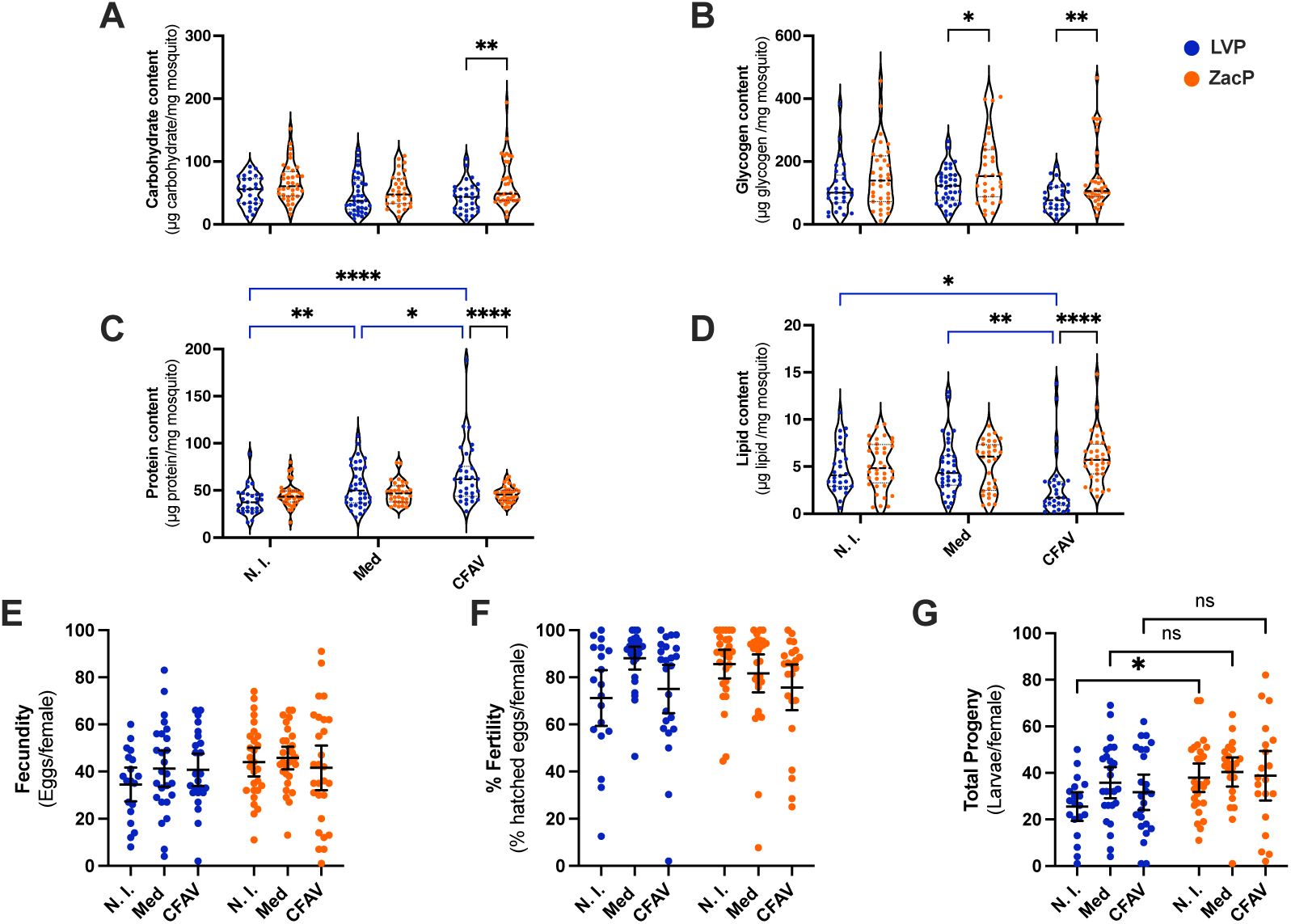
LVP metabolic homeostasis is disrupted after CFAV infection. We quantified the (**A)** carbohydrate content, (**B)** glycogen content, (**C)** protein content and (**D)** lipid content of non-injected (N.I.), media injected (Med) or mosquitoes infected with 10^1.32^ viral particles of CFAV (CFAV), 14 dpi. Each point represents a single mosquito, median is depicted as a dashed lined, blue lines represent comparisons between conditions while black lines represent comparisons between populations. (**E)** Fecundity, **(F)** fertility and **(G)** total progeny of LVP (blue) and ZacP (orange) mosquitoes. Each point represents data from a single mosquito. Median and 95% confidence intervals are shown as lines. ns stands for not significant, * P-value<0.05, ** P-value<0.01, *** P-value<0.001 and **** P-value<0.0001

We detected no differences between populations in any of tested macromolecules in the absence of infection (Fig 4A-D). After CFAV infection, we observed a significant increase in protein content and a decrease in lipid content in LVP mosquitoes (Fig 4 C-D). In contrast, we did not observe significant changes in content of proteins, glycogen and lipids after CFAV infection of ZacP mosquitoes. These results indicate that ZacP maintain metabolic homeostasis after CFAV infection differently than LVP ones.

Finally, we compared the reproductive output of CFAV-infected LVP and ZacP mosquitoes given that viral infections can impair host reproductive capacity [43–45]. We found that fecundity, fertility and total progeny per female were not affected by CFAV infection in either population (Fig 4 E-G).

Altogether these results demonstrate that population-specific management of metabolic homeostasis contributes to modulating *Ae. aegypti* tolerance to a persistent infection and that CFAV infection has a cost for *Ae. aegypti* that manifests in reduced longevity, but not alteration of its reproductive capacity.

## Discussion

The establishment of an infection is a complex phenotype shaped by genetic factors of both host and virus, host physiological status at the time of infection and environmental variables [19, 46, 47]. This complexity leads to infections that can vary from being acute to lifelong persistent depending on how long an organism remains infected. To reduce the burden of an infection hosts have evolved two different, but synergistic, defense strategies called resistance and tolerance. Resistance mechanisms directly target the virus, potentially leading to the clearance of the infection. Tolerance mechanisms reduce detrimental effects of an infection without affecting viral load [48]. Here we show that CFAV establishes a lifelong persistent infection in *Ae. aegypti* mosquitoes, which activates both resistance and tolerance mechanisms. We further show differences in coping with viral infection in LVP and ZacP mosquitoes, with LVP mosquitoes showing a lower tolerance than ZacP mosquitoes. LVP lower tolerance was accompanied by a disruption of metabolic homeostasis.

### Why and how should tolerance and resistance be disentangled?

No infection is fought entirely using resistance or tolerance mechanisms, but the predominance of one mechanism over the other has ecological and evolutionary consequences [46]. An increase in resistance causes a decrease in the prevalence of infection or in viral load, resulting in a strong selective pressure on the virus, which could lead to the emergence of escape mutants [49]. This can result in counter-adaptation by mosquito immunity [50]. A long-term effect of this adaptation and counter-adaptation processes is the fact that immunity genes are known to be among the most fast-evolving mosquito genes [51]. On the other hand, tolerance is not directed to the virus, thus leading to neutral or positive effects in prevalence [52]. The difference between resistance and tolerance has critical consequences in relation to control strategies against arboviruses [53]. A control strategy that focuses on increasing mosquito resistance could apply selective pressure on viruses leading to an ecological unsustainable control [11, 54]. A control strategy that aims at reducing tolerance to viral infections, would lead to an ecologically sustainable approach that targets the elimination of infected mosquitoes [55]. To achieve this goal, molecular mechanisms that control mosquito tolerance to viral infections need to be understood.

Population level differences in viral resistance have been exploited to identify causative genetic loci [56]. A similar approach could be used to uncover genes regulating tolerance. To do this, a key aspect is how to measurement tolerance [20]. Since tolerance has been recognized as in important immunity strategy for animals [57], different quantification methods of tolerance have been proposed [20, 26, 48, 58]. In the first studies of tolerance, tolerance was defined as a reaction norm, which is measured in organisms after infection with different pathogen loads [22, 26, 48]. This strategy allowed to uncover the first example of genetic variation for tolerance in animals [57]. Different mice strains were infected with increasing concentrations of *Plasmodium chabaudi* and mice longevity differed among strains. This result shows that the deterioration of health varies across strains meaning that the level of tolerance differed across strains [57]. An inherent attribute of studying tolerance through a reaction norm is the fact that tolerance is defined as a relative and not absolute measurement [22].

In the case of pathogens resulting in infections with high mortality during their acute stage, two novel ways of assessing tolerance were developed. One method was developed using *Dr. melanogaster*, where multiple genetically different strains are available [20]. The idea is to take advantage of the differences across strains and infect these strains with a single pathogen dose to build a tolerance curve by plotting pathogen burden against the proportion of flies alive two dpi for each strain. Tolerant strains are than identified based on their deviation from the median proportion of alive flies at a given pathogen burden [20]. This approach is constrained by the requirement for a wide variety of strains [20]. A second method also uses a single infection dose and measures the ratio between the VLUD and the likelihood of survival of the infected group relative to a reference group [58]. This strategy has the advantage of resulting in a number, that is an absolute quantification, making it easier to compare tolerance across different hosts or pathogens, but has always been used in scenarios where pathogens reach a plateau load that is lower than the VLUD [58].

CFAV is not a virulent virus for *Ae. aegypti*; infection reaches a plateau that is not significantly different from the VLUD and leads to a decrease in mosquito survival only in older mosquitoes. Given this infection dynamic, a reaction norm analysis is better suited to identify possible subtle differences across populations and/or conditions [25, 26]. Indeed, applying this strategy, we noticed differences in tolerance between LVP and ZacP mosquitos. Using this strategy, we also previously demonstrated differences in tolerance resulting from abiotic stimuli in *Ae. albopictus* [19]. By further combining dose-response curve analyses with viral load quantification at the time of host death, we were able to verify that the decline in host health was not viral-load dependent, implying that tolerance mechanisms contribute to the management of viral infection. The mosquito – virus interaction model that we used in this study serves an ideal platform for further exploring the genetic mechanisms behind *Ae. aegypti* tolerance.

### What are the mechanisms involved in the establishment of *Ae. aegypti* persistent viral infections?

When in 1926 it was proven that *Ae. aegypti* is the vector of DENV, it was suggested that once the infection is acquired, mosquitoes remained infected until their death [59]. This anecdotal conclusion rooted itself in the vector biology community, but, to the best of our knowledge, has never been conclusively proven. Infection of mosquitoes with ISVs are also believed to be lifelong, but, prior to our investigation, analysis were conducted only in live mosquitoes [40]. The fact that all our mosquitoes infected with CFAV had viral genomes at the time of dead unequivocable shows that CAFV produces a systemic lifelong infection.

The persistent infection of CFAV in *Ae. aegpyti* differs from canonical persistent infections of vertebrates in terms of how immunity is modulated after infection, but not in relation to the replication strategies of virus nor the regulation of cellular homeostasis [1]. While vertebrate immunity is evaded by viruses to establish persistent viral infections, *Ae. aegypti* immunity is activated. The production of siRNA molecules evenly covering the CFAV genome indicates an active RNAi response, which regulates viral replication. RNAi activation has been shown crucial in live mosquitoes and in mosquito cell lines to control viral infection [15, 16, 60]. Disruption of the RNAi machinery results in higher viral load, accompanied by increased mortality and cytopathic effects in both mosquitoes and cells [15, 16, 60]. While CFAV reaches a plateau 3-7 dpi, viral load varies among tissues. In both LVP and ZacP mosquitoes, we observed lower viral load in ovaries than carcasses. We further observed that CAFV load in ovaries, but not carcasses, correlates the presence of EVE2 and EVE3 and nrEVEs-derived piRNAs, as previously observed for EVE2 in mosquitoes from Thailand [12]. We hypothesized that the observed reduction in viral load in the ovaries might correlate with the frequency of CFAV vertical transmission or an impact on the reproductive output of mosquitoes after CFAV infection [12, 61]. Accordingly, we observed vertical transmission only in LVP mosquitoes, albeit at a minimal frequency (1.59%). However, we did not observe impairment in the reproductive capacity of either LVP or ZacP mosquitoes following CFAV infection.

During persistent infections in vertebrates, viruses may engage in alternative replication strategies to maintain a non-lytic infection [1]. A recent analysis of the infection dynamic of the ISV Binjari virus in C6/36 cells, a cell line defective for the RNAi response [60], showed that there was an initial acute infection characterize by high viral titer, delayed cell growth and cytopathic effects. This was followed by a persistence infection where the viral titers decreased by 10^5^ fold and stabilized [60]. This phase continued for at least 50 more passages with no pathology and normal cell growth rate [60]. The observation of a shift from acute to persistent infection in a cell line characterized by a disrupted immune response suggests Binjari virus might employ different replication strategies according to the phase of infection. Whether this result translates to other ISVs during persistent viral infections in mosquitoes requires further investigations.

In vertebrate, persistent viral infections may result in homeostasis dysregulation [62]. We also observed a disrupted metabolic homeostasis in LVP mosquitoes, which are less CFAV tolerant than ZacP ones. This result also indirectly emphasizes the role of tolerance mechanisms in managing CFAV infection in mosquitoes. Specifically, we observed a marked reduction in lipid reserves, the primary form of energy storage in mosquitoes [63]. Lipid metabolism is modulated in mosquitoes upon immune challenge [64], but whether this regulation varies across populations and how it influences survival outcomes remains unexplored. In mammals, precise control of mobilization of lipids affects tolerance. For example, production of the growth and differentiation factor 15 (GDF15) after bacterial infection in mice enables the liver to export triglycerides; blocking GDF15 with a monoclonal antibody leads to impaired cardiac function and increases mortality without affecting pathogen load [65]. Replication of DENV, West Nile Virus and Japanese encephalitis virus occurs near the endoplasmic reticulum (ER), inducing ER rearrangement, proliferation and dilation [66]. ER stress can lead to an increase in miss-folded or unfolded proteins triggering unfolded protein response (UPR) [67]. UPR activation increases ER capacity to maintain ER homeostasis [68]. Alterations of the proper regulation of this pathway after infection could lead to increased energetic needs and ultimately to a decreased survival. Our observation of an increase in protein content after CFAV infection of LVP mosquitoes is indirect evidence of this hypothesis.

## Materials and methods

### Mosquito Rearing

We used the Liverpool (LVP) and Zac Panda (ZacP) laboratory populations and maintained all mosquitoes in climatic chambers (Binder KBWF) at a constant 28 °C temperature, 70 ± 5% relative humidity, and a 12:12-hour light/dark photoperiod. We reared 200 larvae in 19 × 19 × 16 cm plastic pans (BudDorm) filled with one liter of distilled water and added ten Tetra Goldfish Gold Colour fish food pellets (Tetra Werks, Germany) daily. After pupation, we transferred mosquitoes to BugDorm cages (30 cm²) and provided them with a 20% sugar solution *ad libitum*. Seven to ten days after adult emergence, we offered defibrinated mutton blood (Biolife Italiana) using a Hemotek feeding apparatus.

### Viral Preparation

We obtained the CFAV (Rio Piedras 2002, Ref-SKU: 001v-EVA68) from Ronald Van Rij (Radboud University, Nijmegen, NL). We propagated the virus in C6/36 cells maintained in L-15 Complete Medium (Leibovitz’s L-15 Medium; Gibco, Ref:11415-049) supplemented with 10% FBS (Gibco, Ref:16000-044 Lot: 2421078RP), 2% TBP (Gibco, 18050-039), 1× MEM NEAA (Gibco, 11140-035), and 2% Penicillin/Streptomycin (Sigma-Aldrich, P0781-100 mL).

We seeded two T75 tissue culture flasks with identical volumes of cells and allowed them to reach 80% confluency. We used one flask to count the cells and infected the other with a multiplicity of infection (MOI) of 0.1 (i.e., one viral particle per ten cells). We let the virus replicate for five days. After incubation, we collected the medium, clarified it by centrifugation at 4000 × *g* for ten minutes at 4 °C, and concentrated it using an AMICON Ultra-15 100k centrifugal filter (Millipore, UFC910008) following the manufacturer’s instructions. We quantified the viral stock by a focus forming assay and stored it at –80 °C until use.

### Viral Titre Quantification

We seeded 0.75 × 10⁵ C6/36 cells in each well of a 96-well plate and incubated them overnight at 28 °C. The next day, concentrated virus and three tenfold serial dilutions were added to the wells and incubated at 28 °C for one hour with shaking at 180 rpm. After incubation, the medium was removed, and 100 µL of a 1:1 mixture of L-15 complete medium and 3.5% carboxymethylcellulose sodium salt (medium viscosity, Sigma, C488-500G) in water was added to each well.

At 96 hours post-infection, the overlay was removed, and wells were washed with 100 µL of PBS (Medicago, 12-9423-5). Then, 100 µL of an 80:20 acetone:PBS solution was added, and the plate was incubated at −20 °C for 20 minutes. The acetone was then discarded, and the plate was allowed to dry overnight. The fixed plate was stored for up to one week before visualization.

On the day of microscopy, each well was washed twice with PBST (PBS containing 0.05% Tween 20; Sigma, P1379-500 mL). Then, 50 µL of 3% blocking reagent (Cytbia, RPN2125) in PBST was added, and the plate was incubated at 37 °C for 30 minutes with shaking at 180 rpm. After blocking, the solution was discarded, and 50 µL of a 1:1000 dilution of primary antibody (mouse anti-CFAV envelope, GenScript) in PBST was added. The plate was incubated for two hours at 37 °C with shaking at 180 rpm.

Following incubation, the primary antibody was discarded, and wells were washed three times with PBST. Then, 50 µL of a 1:2000 dilution of secondary antibody (goat anti-mouse Alexa Fluor Plus 647) in PBST was added, and the plate was incubated for another two hours at 37 °C with shaking. Finally, the secondary antibody was removed, wells were washed three times with PBST, refilled with 100 µL of PBS, and images were acquired using a Leica DMi8 microscope with an HC PL FLUPTAR 4x objective. Each sample was seeded twice and the counted particles were averaged.

### Viral Infection

We used three- to eight-day-old females for all infection experiments. We always initiated infections seven hours after the start of the light cycle to reduce circadian cycle bias [69]. We cold-anesthetized the mosquitoes and divided them into three groups: non-injected (N.I.), media-injected (Med), and CFAV-infected (CFAV). We injected 50 nL of either CFAV or L-15 complete medium intrathoracically using a Nanoject III (Drummond). We kept the injection volume constant and varied the virus concentration for tolerance assays. After infection, we housed up to 20 females per group in cardboard cups (0.266 m²) under standard rearing conditions.

### Viral Load Quantification

We extracted RNA from either whole mosquitoes or individual tissues at specific timepoints post-infection. We used 500 µL TRIzol Reagent (Invitrogen, Ref: 15596018) per mosquito or tissue and followed the manufacturer’s protocol. We resuspended the final RNA pellet in 20 µL of ultra-pure water. We used 10 µL of the RNA to synthesize cDNA with the GoScript Reverse Transcription Mix (Promega), following the manufacturer’s instructions. We performed qPCR using the QuantiNova SYBR Green PCR kit (Qiagen) with primers targeting the *ns3* viral gene [12] and the *rps17* housekeeping gene [70]. We calculated relative viral loads using the 2^−ΔΔCt^ method.

### Tolerance

We assessed tolerance by measuring survival after mock infection or infection with five viral doses: 10^1.32^, 10^2.32^, 10^3.32^, 10^4.32^, and 10^5.32^ CFAV particles. For each dose, we used 20 mosquitoes and repeated the experiment three times. We excluded mosquitoes that died within three days post-injection from analysis, assuming their deaths were caused by injury during injection.

### Gene Expression

We performed RNA extraction, cDNA synthesis, and qPCR as described in the “**Viral Load Quantification”** section. For gene expression analysis, we used primers for *dcr2* and *ago2* as genes of interest and *rps17* as the housekeeping gene (Supplementary Table1).

### sRNA Sequencing

Small RNA libraries were generated from six whole mosquitoes or ovary pairs from LVP and ZacP mosquitoes. Total RNA was extracted and sent to BGI for sRNA sequencing, trimmed reads datasets were provided and processed as follow. Trimmed reads were generated by removing adapters, low-quality bases (Phred score<20), reads enriched with a poly(A) tail, and reads with length <15 or >44 basepairs with SOAPnuke [71] using the options “-n 0.001 -l 20 -q 0.1 --highA 1 --minReadLen 15 --maxReadLen 44 - ada_trim”. After a quality control (QC) analysis of the trimmed reads performed with FASTQC v0.11.9 [72], reads were aligned separately to three reference sequences, the Cell Fusion Agent virus (CFAV) genome strain Rio Piedras02 (GenBank accession number GQ165810), the endogenous viral element (EVE) CFAV-EVE-3 from Crava *et al.* (2021) and the EVE2 from Suziki et al. (2020) using Bowtie v1.3.1 [12, 41, 73]. Aligned reads were sorted and processed to create sequence alignment map (SAM) files from binary alignment map (BAM) files with SAMtools v1.10 [74]. An extensive QC report of trimmed reads and alignments for all samples was prepared using MultiQC v.1.13 [75], and it is available at the GitHub repository: https://github.com/naborlozada/Perdomo_et_al_2025_sRNA_analysis.

The distribution and quantification of small RNAs across both reference sequences were estimated separately with a custom R script using the R packages tidyverse v1.3.1 [76] and Biostrings v2.54.0 [77]. Briefly, information of reads counts, sequence length, mapping strands, and genomic position of small RNAs were extracted from SAM files, then small RNAs count frequencies mapped either to one or both strands were calculated only for sequence lengths ≥18 bp and ≤33 bp. The small RNAs distribution across each reference sequence for both strands was obtained separately for small interfering RNA (21 bp) and Piwi-interacting RNAs (26-to-30 bp). Small RNAs frequencies and distributions for both target sequences were plotted with Graphpad Prism 10.4.0.

### EVE Presence

We extracted DNA from legs and wings using the Monarch Genomic DNA Purification Kit (#T3010L), following the animal tissue protocol and incubating the samples for 60 minutes to ensure tissue lysis. We resuspended the DNA in 35 µL of ultra-pure water. We performed PCR with *Ae. aegypti* specific 18S primers to confirm successful DNA extraction, followed by two PCRs to detect EVE3 (Crava) and EVE2 (Suzuki) [12, 41].

### Energy Reserves Quantification

We assessed energy reserves, including carbohydrates, glycogen, protein, and lipid content, in individual mosquitoes 14 dpi using a modified Foray et al. protocol, as described in Carlassara et al. [78, 79].

We weighed each mosquito using a microbalance (Mettler AC100) and homogenized them in 180 µL of lysis buffer (100 mM KH₂PO₄, 1 mM DTT, 1 mM EDTA, pH 7.4). We centrifuged the samples at 180 × *g* for 5 minutes at 4 °C (Eppendorf 5804R) and used the supernatant for subsequent assays. For protein quantification, we mixed 2.5 µL of supernatant with Bradford reagent (Sigma-Aldrich, B6916) and measured absorbance at 595 nm using a CLARIOstar plate reader. We used BSA as the standard.

To extract carbohydrates and lipids, we added 20 μL of 20% Na₂SO₄ and 1.5 mL of a chloroform–methanol (1:2) solution to the remaining supernatant, then centrifuged at 180 × *g* for 15 minutes at 4 °C. We set aside 100 μL of the supernatant for lipid quantification and used the remainder for carbohydrate analysis. We reacted the latter with anthrone reagent (1.42 g L⁻¹ in 70% H₂SO₄) at 90 °C for 15 minutes and measured absorbance at 625 nm.

We washed the remaining pellet twice with 80% methanol and reacted it with anthrone reagent to determine glycogen content. For both carbohydrate and glycogen assays, we used D-glucose as the standard.

To quantify lipids, we heated the 100 μL supernatant to 90 °C, treated it with 10 μL of H₂SO₄, and incubated it for 2 minutes. We then cooled the sample, reacted it with 190 μL of vanillin reagent (1.2 g L⁻¹ in 68% H₃PO₄), and measured absorbance at 525 nm.

### Reproductive Capacity

We allowed mosquitoes one week to recover after injection. At 7 days post-injection, we offered a blood meal to mosquitoes from the N.I., Med, and CFAV groups and followed Tsujimoto and Adelman protocol detailed in the Journal of Visualized Experiments [80].

### Statistics and Reproducibility

We performed all figures and statistical analyses using GraphPad Prism version 10.4.0. S1 Table reports all statistical results.

### Data Availability Statement

We deposited all data supporting the findings of this study in Zenodo at https://zenodo. All trimmed read datasets are available in the NCBI Sequence Read Archive under BioProject accession code: **PRJNA1287247**.

## Acknowlodgments

We are grateful to Lino Ometto of the Univesrity of Pavia for his input in the manuscript, Brian Schaub and all members of the Bonizzoni Lab for fruitful discussions. We would like to thank Amanda Oldani of Centro Grandi Strumenti from the Microscopia Ottica facility for her support and assistance in this work.

## Supporting Information

S1 Fig. Vertical transmission

S2 Fig. sRNA sequencing ovaries of LVP mosquitoes

S3 Fig. sRNA sequencing mapping to EVE2 and EVE3

S1 Table. Statistical analysis performed in this work

S2 Table. Primer table

